# Developmental system drift in motor ganglion patterning between distantly related tunicates

**DOI:** 10.1101/320804

**Authors:** Elijah K. Lowe, Alberto Stolfi

## Abstract

The larval nervous system of the solitary tunicate *Ciona* is a simple model for the study of chordate neurodevelopment. The development and connectivity of the *Ciona* Motor Ganglion (MG) has been studied in fine detail, but how this important structure develops in other tunicates is not well known. By comparing gene expression patterns in the developing MG of the distantly related tunicate *Molgula occidentalis*, we found that its patterning is highly conserved compared to the *Ciona* MG. MG neuronal subtypes in *Molgula* were specified in the exact same positions as in *Ciona*, though the timing of subtype-specific gene expression onset was slightly shifted to begin earlier, relative to mitotic exit and differentiation. In transgenic *Molgula* embryos electroporated with *Dmbx* reporter plasmids, we were also able to characterize the morphology of the lone pair of descending decussating neurons (ddNs) in *Molgula*, revealing the same unique contralateral projection seen in *Ciona* ddNs and their putative vertebrate homologs the Mauthner cells. Although *Dmbx* expression labels the ddNs in both species, cross-species transgenic assays revealed significant changes to the *cis-*regulatory logic underlying *Dmbx* transcription. We found that *Dmbx cis-*regulatory DNAs from *Ciona* can drive highly specific reporter gene expression in *Molgula* ddNs, but *Molgula* sequences are not active in *Ciona* ddNs. This acute divergence in the molecular mechanisms that underlie otherwise functionally conserved *cis*-regulatory DNAs supports the recently proposed idea that the extreme genetic plasticity observed in tunicates may be attributed to the extreme *rigidity* of the spatial organization of their embryonic cell lineages.

## Introduction

Tunicates have a long history as tractable laboratory organisms for the study of embryonic development (Satoh, 2013). Most tunicate larvae develop rapidly but invariantly, according to highly stereotyped cell lineages. Furthermore, many also possess highly compact genomes and are quite amenable to a wide variety of molecular assays and perturbations. More recently, tunicates have also begun emerging as model organisms for developmental neurobiology (Nishino, 2018). The complete connectome of the larva of the tunicate *Ciona intestinalis*, the second connectome ever mapped after that of the nematode *C. elegans* (White et al., 1986), revealed the synaptic connections of all 177 neurons of the central nervous system (CNS) and all 54 neurons of the peripheral nervous system (PNS)(Ryan et al., 2016, 2018). With 231 total neurons (CNS and PNS combined), the *Ciona* larval nervous system is one of the smallest ever described, smaller than even the nervous system of the *C. elegans* hermaphrodite (302 neurons).

The *Ciona* connectome further revealed specific neural circuits that are conserved between tunicates and their sister group, the vertebrates, including a putative homolog of the Mauthner cell/C-start escape response circuit of fish (Ryan et al., 2017). In *Ciona*, the putative Mauthner cell (M-cell) homologs are a single pair of descending decussating neurons (ddNs), which correspond to the A12.239 pair of cells of the Motor Ganglion (MG), a cluster of 30 neurons that comprise a central pattern generator for the swimming behavior of the larva (Nishino et al., 2010), and proposed to be homologous to a combination of the vertebrate hind brain and spinal cord (Wada et al., 1998). The development of the ddNs and the rest of the MG has been studied in some detail, revealing gene expression patterns and transcriptional regulatory networks that are shared with hindbrain and spinal cord development in vertebrates (Imai et al., 2009; Stolfi and Levine, 2011; Stolfi et al., 2011). These close parallels are especially striking considering the obvious, drastic reduction in size and complexity of the *Ciona* MG relative to the corresponding regions in vertebrates. However, it was found that cell fate specification and transcriptional patterning in the *Ciona* MG depends largely on cell-cell contact-dependent signaling within the neural tube by the Delta/Notch and Ephrin/Eph pathways, not on gradients of secreted, long-range morphogens as in the vertebrate spinal cord (Hudson et al., 2011; Stolfi et al., 2011). Tunicates have also garnered recent attention for the fact that their extremely reduced, stereotyped cell lineages are highly conserved even between distantly related species. In spite of highly elevated mutation rates genome sequence divergence, and deep evolutionary timescales (Delsuc et al., 2018; Kocot et al., 2018; Tsagkogeorga et al., 2012; Tsagkogeorga et al., 2010), the embryos of distantly related solitary tunicates like *Ciona* and *Molgula* (estimated divergence 390 million years apart)(Delsuc et al., 2018), are nearly indistinguishable (Stolfi et al., 2014). We previously compared the development of the cardiopharyngeal mesoderm between *Ciona robusta* and *Molgula occidentalis*, and found that there were virtually no differences in cell lineage and gene expression. However, we did find that underneath this seemingly conserved developmental program, there were considerable cryptic functional differences, resulting in *cis*-regulatory “unintelligibility” between *Ciona* and *Molgula.* In other words, homologous *cis*-regulatory elements driving identical gene expression patterns were partially or completely non-functional in cross-species transgenic assays. This phenomenon was ascribed to a particularly acute form of developmental system drift (DSD)(True and Haag, 2001). It was recently proposed that the acute DSD observed in solitary tunicate evolution may be directly related to their mode of development, which depends primarily on invariant, contact-dependent intercellular signaling for cell fate induction. This geometric constraint would have relaxed constraints on genome evolution imposed by the relatively intricate and immutable transcriptional networks and *cis-*regulatory logics required for inductive events in species with larger, more variable embryos (i.e. the “Geometry vs. Genes” paradigm)(Guignard et al., 2018).

Here we report that the phenomenon of acute DSD also extends to neurodevelopment in relatively late phases of tunicate embryogenesis. By surveying the development of the *M. occidentalis* MG, we show that transcription factor expression patterns, or “codes”, in the developing MG are identical between *Molgula* and *Ciona*, indicating near perfect conservation of arrangement of neuronal subtypes. However, cross-species reporter assays revealed acute DSD in the regulation of *Dmbx* transcription in homologous ddN precursor cells. Our results support a “Geometry vs. Genes” model to explain the emergence of DSD in the tunicate MG, given its patterning by invariant cell-cell contacts and the parsimony of its final configuration as inferred by the *Ciona* connectome.

## Materials and methods

### *Molgula occidentalis* genome assembly improvement

Libraries prepared for the version of the *Mogula occidentalis* genome (ELv1.2) currently available on ANISEED (https://www.aniseed.cnrs.fr/)(Brozovic et al., 2017; Brozovic et al., 2015; Stolfi et al., 2014) were reused to reassemble the genome. The original 3 DNA libraries of paired-end reads had insert sizes of 300–400, 650–750, and 875–975 base pairs (NCBI SRA ID# PRJNA253689)(Stolfi et al., 2014). Reads were first filtered to a kmer coverage of 100X using the Khmer suite (Crusoe et al., 2015). Assembly was done using Oases (v 0.2.08)(Schulz et al., 2012) at various word lengths (k), with k = 67 being selected as the assembly to continue the downstream analyses. After reassembly (version “k67”), additional scaffolding was done using Redundans (Pryszcz and Gabaldón, 2016) producing the version “k67_R” (available at https://osf.io/3crup/). The ANISEED genome (ELv1.2) was scaffolded with Redundans as well, for comparison (version “ANISEED_R”). First redundant contigs were detected and selectively removed, next genome fragments were joined using the previously mentioned libraries from the initial assembly. Finally, gapped regions of the scaffold were filled using these 3 paired-end libraries. After reassembly and additional scaffolding, the scaffolding quality was examined using REAPR (v1.0.17)(Hunt et al., 2013) mapping all three libraries to back to the assembly.

### *Molgula occidentalis* transcriptome sequencing

Gravid *Molgula occidentalis* Traustedt, 1883 adults were collected and shipped by Gulf Specimen Marine Lab (Panacea, FL). Eggs were fertilized as previously described (Stolfi et al., 2014). *M. occidentalis* embryos and larvae were grown at ~24°C and collected into three separate stage pools: 0.33 hours post-fertilization (hpf), 7 hpf, and 16 hpf. Total RNA was extracted from each pooled sample using RNAqueous Total RNA Isolation kit (ThermoFisher). PolyA+ mRNAs were selected using Oligo d(T)25 magnetic beads (New England Biolabs) following two rounds of the manufacturer’s protocol. Directional RNAseq libraries were prepared according to a modified version of the protocol contained at: https://wikis.nyu.edu/pages/viewpage.action?pageId=24445095 (Neymotin et al., 2014). First strand synthesis was performed using Super Script III (ThermoFisher), primed with Oligo d(T) and random hexamer primers. Second-strand synthesis was performed using dUTP instead of dTTP for directional (strand-specific) sequencing (Parkhomchuk et al., 2009). Samples were processed and ligated to NETFLEX DNA barcode adapters (BioO Scientific) for multiplex sequencing. Adapter dimers were removed using AMPure beads (Agencourt). Samples were then treated with Uracil-DNA Glycosylase, amplified using 12 cycles of PCR, and purified once with a minElute kit (Qiagen) and once with AMPure beads. Samples were sequenced by Illumina 2000 2x50 bp runs on two lanes (all three samples were multiplex-sequenced in each lane). Resulting sequences will be deposited in the NCBI SRA database (accession number pending).

### Transcriptome assembly and gene model improvement

Paired-end sequences generated above were used to generate gene model. Reads were quality filtered/trimmed using trimmomatic (v0.33)(Bolger et al., 2014) and the following parameters “MINLEN:25, and sliding window of 4 with a minimum score of 5 (ILLUMINACLIP:TruSeq3-PE.fa:2:30:10 SLIDINGWINDOW:4:5 MINLEN:25)”. Each of three embryonic stage libraries (0.33 hpf, 7 hpf and 16 hpf) were individually mapped to the various genome versions using hisat2 (v 2.1.0)(Kim et al., 2015), with the default parameters. The sam files generated from hisat2 were then sorted and converted to bam using samtools (v 1.5)(Li et al., 2009) and merged using PicardTools with java version 1.8 (http://broadinstitute.github.io/picard/). The merged bam file was then processed using StringTie (v 1.3.3b)(Pertea et al., 2015) to create gene models. Gene models were then extracted using the gffread utility from cufflinks (Trapnell et al., 2012) and evaluated using BUSCO for completeness (Simão et al., 2015). Transcripts/gene models have been deposited at OSF (https://osf.io/3crup/).

### Fertilization, dechorionation, and electroporation of embryos

*M. occidentalis* eggs were fertilized, dechorionated, and/or electroporated as previously described (Stolfi et al., 2014). *Ciona robusta* (*intestinalis* Type A) adults were collected and shipped by M-REP (San Diego, CA) and eggs dechorionated, fertilized, and electroporated as previously described (Christiaen et al., 2009b). *Dmbx cis-*regulatory DNAs from two *Molgula* species (*M. occidentalis* and *M. oculata*) were cloned intro reporter plasmids as illustrated in supplemental sequences files (**Supplemental File 1**). *Cirobu.Dmbx>Unc-76::Venus* and *Cirobu.Fgf8/17/18>H2B::mCherry* reporter plasmids were published previously (Stolfi and Levine, 2011; Stolfi et al., 2011).

### mRNA probe synthesis and *In situ* hybridization

Templates for antisense riboprobes for *in situ* hybridization were amplified by PCR or SMARTer 3’/5’-RACE (Clontech) from cDNA libraries or from genomic DNA (see **Supplemental File 1** for details on each sequence). Template sequences were cloned either using TOPO-TA cloning (ThermoFisher) into pCRII dual promoter vector, or using restriction enzyme cloning into pCiProbe (see **Supplemental File 1**) linearized NotI-EcoRI. *In vitro* transcription of labeled riboprobes and two-color fluorescent *in situ* hybridization were performed as previously described (Ikuta and Saiga, 2007b).

## Results and discussion

### *M. occidentalis* transcriptome assembly and gene prediction

We recently sequenced the genomes of 3 species in the genus *Molgula*: *M. occidentalis, M. oculata*, and *M. occulta* (Stolfi et al., 2014), which can be browsed freely on the Tunicate molecular biology database ANISEED (https://www.aniseed.cnrs.fr/)(Brozovic et al., 2017). Of these, *M. occidentalis* emerged as a valuable species for comparative studies of tunicate development, mainly because their zygotes can be transfected with plasmid DNAs via electroporation, much like the major tunicate laboratory model species in the *Ciona* genus (*C. intestinalis, C. robusta*, and *C. savignyi)*. To help establish additional molecular tools for developmental studies in *M. occidentalis*, we assembled a transcriptome based on RNAseq of 3 stages of embryonic development. These were then used to predict a new set of gene models found in the *M. occidentalis* genome sequence.

To do this, we also reassessed our previous *M. occidentalis* genome assemblies. In previous assemblies, some of the low-coverage regions were removed using khmer to reduce assembly fragmentation (Crusoe et al., 2015). To preserve the information contained in these low-coverage regions, we re-assembled the genome from the original raw sequencing reads (see **Materials and methods** for details). While this new assembly initially had an N50 of only 519 bp and over 680,000 scaffolds, additional scaffolding decreased the number of scaffolds to 30,188, with 20,616 of those (68%) being over 1 kb in length (Table 1). As a result of this better scaffolding, the N50 also increased from 519 to 18,312. The gap filling process decreased the number of missing bases (Ns) from 551,0404 to 189,152. The end result is the “k67_R” genome assembly version, which we have deposited at OSF (https://osf.io/3crup/). While these procedures improved the genome assembly statistics, we wanted to ensure that the additional gene content was still preserved. To check this, we mapped mRNA reads and produced gene models to be tested using BUSCO (Simão et al., 2015). This allowed us to compare our new gene models against a defined collection of highly conserved sequences expected to be present in all metazoans, as a measure of assembly completeness. Our BUSCO results (Fig 1) indicated that fewer reference genes were missing (73 missing out of 843 total) in the new “k67_R” assembly than in the ANISEED ELv1.2 version (141/843 missing) or even a version of the ANSEED assembly with additional scaffolding performed (“ANISEED_R”, 79/843 missing). This indicates that these new versions of the genome assembly and their corresponding gene models should enhance the identification and cloning of sequences to use as molecular tools for *M. occidentalis.*

**Figure 1.**
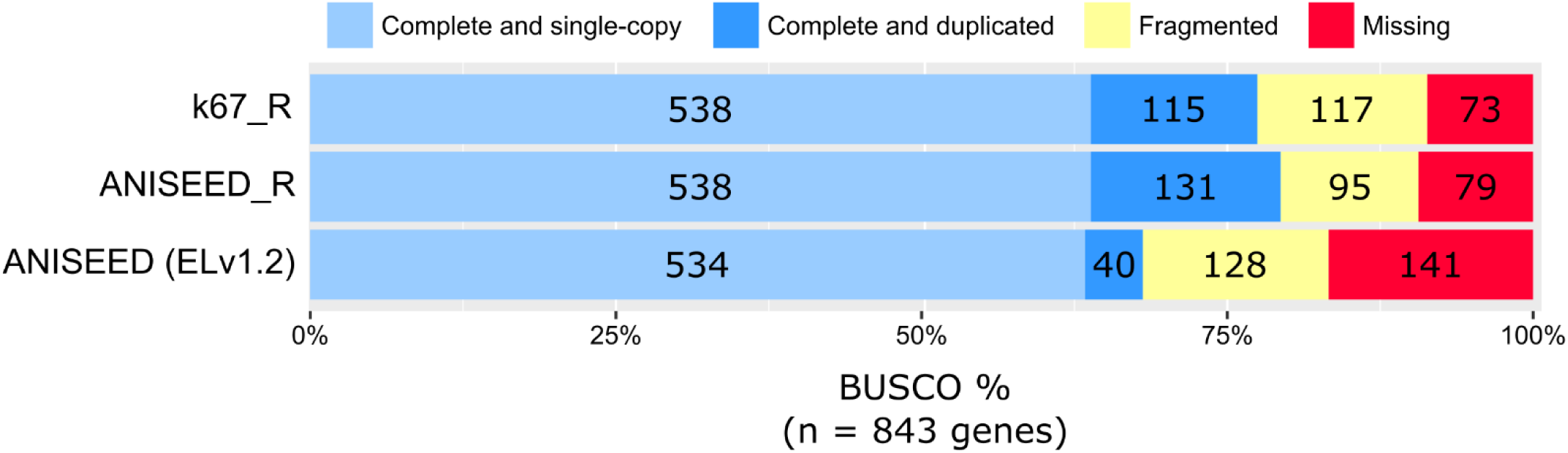
Benchmarking Universal Single-Copy Orthologs (BUSCO) in *M. occidentalis*. BUSCO was used to asses gene model predictions based on the previously published and released *M. occidentalis* genome assembly (ANISEED ELv1.2) and on the re-assemblies from this study (k67_R and ANISEED_R). BUSCO is used as a measure of completeness of gene model sets predicted from a genome assemblies, searching for 843 highly conserved genes that should be present in single copies in >90% of metazoan genomes. There are five categories of gene recovery: complete and single-copy, complete and duplicated, fragmented, and Missing. Complete genes are those that preserve ~95% of the gene length. Single-copy are genes with only one copy in the gene model set and duplicated are those with multiple copies identified either by gene duplication or assembly errors. Fragmented genes are those recovered with less than 95% of the gene length, and missing are those that are not found to be present at all.

**Table 1.** *M. occidentalis* genome reassembly statistics

| Assembly version | k67_R | ANISEED_R | ANISEED* |
| --- | --- | --- | --- |
| Total length (bp) | 207,168,883 | 242,783,314 | 262,547,660 |
| N's | 189,152 | 885,908 | 33,473,590 |
| Scaffolds | 30,188 | 16,148 | 51,761 |
| Scaffolds $\geq 1$ kb | 20,616 | 13,751 | 21,251 |
| Longest scaffold (bp) | 250,303 | 235,023 | 229,829 |
| N50 (bp) | 18,312 | 35,542 | 26,298 |

\* ELv1.2 assembly version www.aniseed.cnrs.fr/fgb2/gbrowse/moocci_elv12/

### Gene expression patterns in the *M. occidentalis* Motor Ganglion

The *Ciona* MG is comprised of 22 neurons and includes a “core” MG of 5 morphologically and molecularly distinct left/right pairs of neurons (Ryan et al., 2016). In the remainder of this study, we will only refer to the neurons on one side, for simplicity. The core MG is derived from the A9.30 and A9.32 blastomeres of the neural plate (Fig 2A) that will give rise to cells along the lateral rows of the neural tube after neurulation and neural tube closure (Cole and Meinertzhagen, 2004; Navarrete and Levine, 2016; Nicol and Meinertzhagen, 1988a, b). At the level of the MG, the neural tube of the *Ciona* embryo is formed by only four single-file rows of cells oriented along the anterior-posterior (A-P) axis: a dorsal row (roof plate), a ventral row (floor plate), and two neurogenic lateral rows from which most of the core MG neurons are specified (Fig 2B). Ascending contralateral inhibitory neurons (ACINs) derived from the A9.29 blastomeres (Nishitsuji et al., 2012) have been traditionally excluded from the MG based on their more posterior location at the base of the tail. However, these are likely indispensable cogs in the MG central pattern generator, driving left/right alternation of tail contractions during swimming by glycinergic neurotransmission (Nishino et al., 2010). Remaining MG neurons are poorly studied and their development is largely unknown. In this study, we focused on the “core” MG neurons, those derived from the A9.30 and A9.32 lineages, because these lineages have been the most thoroughly-studied MG lineages (Imai et al., 2009; Navarrete and Levine, 2016; Stolfi and Levine, 2011). Expression patterns of developmental regulators and provisional gene regulatory networks have been documented in these lineages (Imai et al., 2009), and the induction events responsible for specifying the 5 distinct types of neurons in this core MG have also been elucidated (Stolfi and Levine, 2011; Stolfi et al., 2011). Thus, the MG is a perfect starting point for a comparative study on the evolution of neurodevelopment in tunicates.

**Figure 2.**
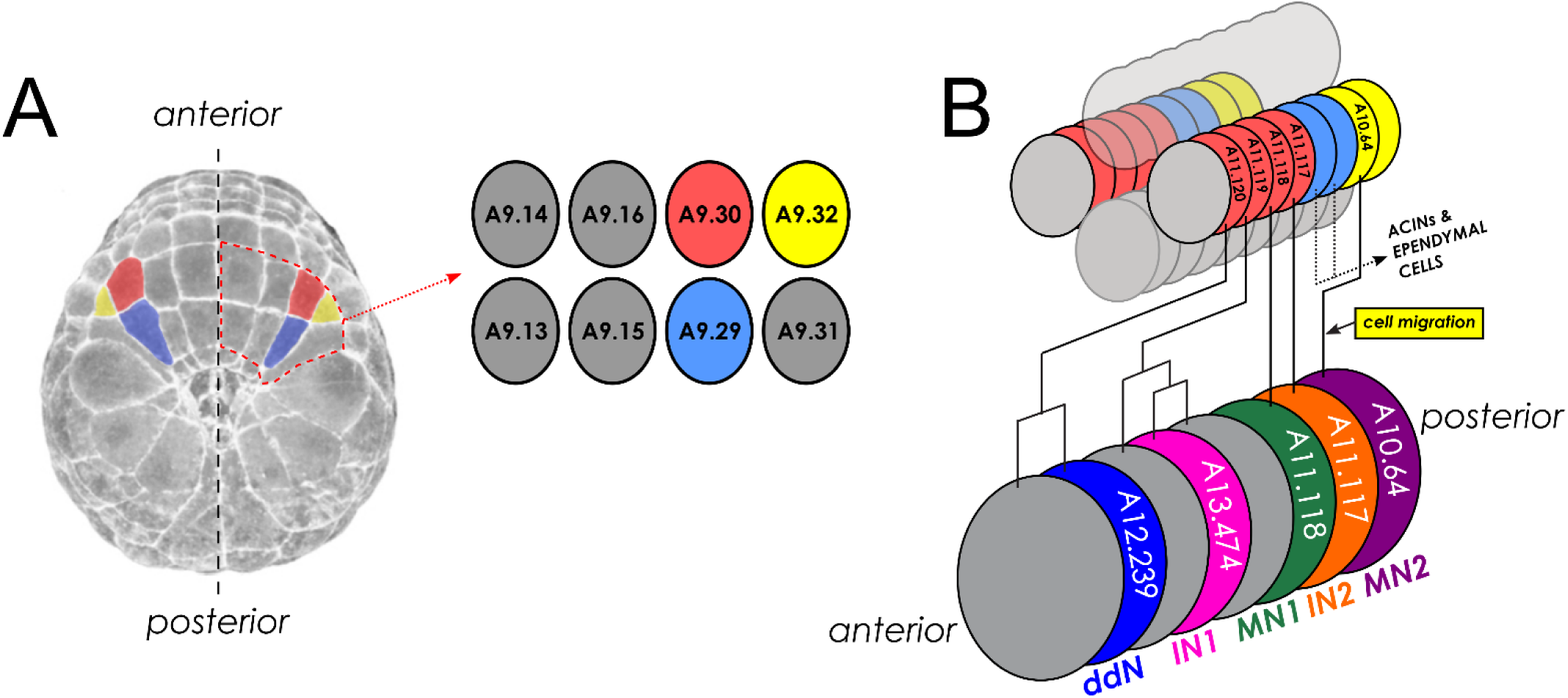
Motor ganglion lineages and neuron subtype configuration in *Ciona*. **A)** Dorsal view of a late gastrula-stage *Ciona robusta* embryo highlighting in false color the bilaterally symmetric right/left pairs of blastomeres in the vegetal neural plate that give rise to the “core” Motor Ganglion and Anterior Caudal Inhibitory Neurons (ACINs) of the nerve cord: A9.30 (red), A9.32 (yellow), and A9.29 (blue). Black dashed line indicated embryo midline. Red dashed outline denotes the right vegetal neural plate annotated in the inset diagram, which indicates the exact cell identities according to the established cell lineage nomenclature (Conklin 1905). Embryo image adapted from the Four-dimensional Ascidian Body Atlas website (FABA, http://chordate.bpni.bio.keio.ac.jp/faba/1.4/top.html)(Hotta et al. 2007) **B)** Diagram representing a section of the dorsal hollow neural tube derived from the neural plate after the process of neurulation. Cells of the neural tube are color-coded according to their lineages using the same color scheme as the neural plate diagram in (A). The neural tube is comprised exactly of four single-file rows of cells: 1 dorsal row, 2 lateral rows, and 1 ventral row. Cells of the lateral rows contributing to the bilaterally symmetric Motor Ganglion are: A11.117-A11.120, derived from A9.30, and A10.64, derived from A9.32. The A9.29 lineage, which gives rise to ACINs and ependymal cells posterior to the MG, is not illustrated in great detail, for the sake of simplicity. At bottom is a diagram of the 5 core MG neuron types. Black lines indicate their descent patterns from those cells at top. ddN = descending decussating neuron, IN1 = interneuron 1, IN2 = interneuron 2, MN1 = motor neuron 1, MN2 = motor neuron 2. Non-neuronal cells are grey. Cells are identical on both left and right sides.

We previously showed that the overall development of the *M. occidentalis* embryo is very similar to the development of the *Ciona* embryo (Stolfi et al., 2014). An in-depth study of the cardiopharyngeal mesoderm (B7.5 cell lineage) revealed perfect conservation of precise cell divisions and fate specification events, with only minimal differences in morphogenesis and timing of gene expression. However, other tissues beyond the cardiopharyngeal mesoderm were not surveyed. In this study, we sought to extend our understanding of tunicate evolution by comparing the developing nervous systems of *M. occidentalis* and *C. robusta* (*intestinalis* Type A). *In situ* mRNA hybridization (ISH) for neural marker *Celf3/4/5* (formerly *Etr-1*) in *M. occidentalis* neurula embryos revealed the CNS developing from the lateral rows of the dorsal hollow neural tube, as in *Ciona* (Satou et al., 2001)(Fig. 3A). Later, at tailbud stages, ISH for the neuronal transcription factors *Neurogenin, Onecut*, and *Ebf* revealed ongoing neuronal specification in the brain, MG, and bipolar tail neurons of the tail (Fig 3B-D), also identical to the expression of these regulators in *Ciona* embryos (Mazet et al., 2005; Satou et al., 2001). Two-color ISH of *Ebf* and the cholinergic marker *Slc18a* (also known as *Vesicular acetylcholine transporter*, or *VAChT*)(Takamura et al., 2002) revealed the earliest differentiating neurons of the larval CNS, the motor neurons of the MG (Fig 3D). Given that no significant differences were revealed between *Molgula* and *Ciona* using these broad markers of neural fate, we focused our attention specifically to the developing MG, where we would be able to analyze more subtype-specific gene expression and cell fates.

**Figure 3.**
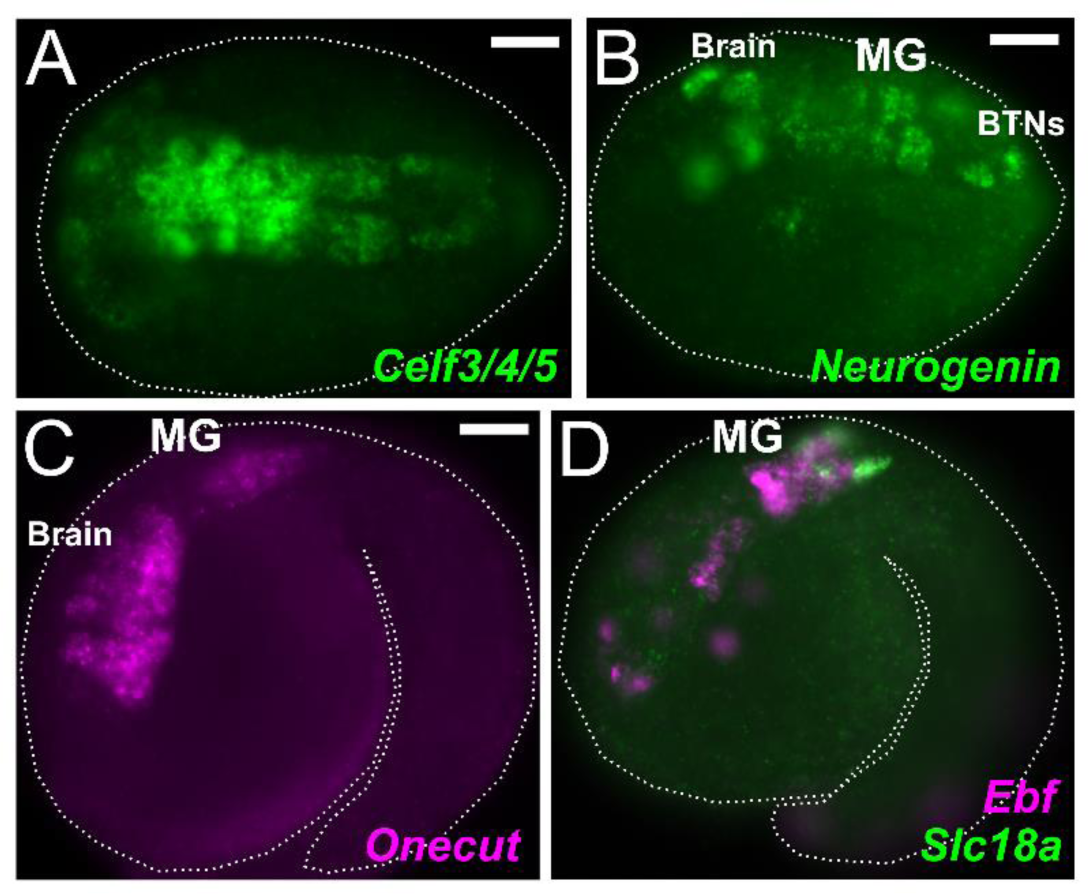
The developing central nervous system of *Molgula occidentalis*. **A)** Dorsal view of a neurula stage *Molgula occidentalis* embryo. Green shows fluorescent *in situ* hybridization for *Celf3/4/5* (also known as *Etr-*1) transcription. **B)** *In situ* hybridization for *Neurogenin* in initial tailbud embryo, showing transcription in the embryonic brain, motor ganglion (MG), and bipolar tail neurons (BTNs). **C)** *In situ* hybridization for *Onecut*, expressed in brain and MG. **C)** Two-color fluorescent *in situ* hybridization showing co-expression of *Ebf* (magenta) and *Slc18a* (also known as *VAChT*, green) in the MG. Embryo outlined by dotted lines. Anterior is to the left in all images. All scale bars = 25 µm.

In *C. robusta*, the expression patterns of mRNAs encoding transcription factors are dynamically regulated in the developing MG (Ikuta and Saiga, 2007a; Imai et al., 2009). Some of these patterns comprise a conserved “code” of homeodomain-containing genes that are differentially expressed along the dorsal-ventral (D-V) developing spinal cord of vertebrate embryos (Stolfi et al., 2011). However, in the developing *Ciona* MG, the neurogenic domain of neural tube is restricted to the single-file lateral rows of cells. Therefore the D-V code of the vertebrate spinal cord is transposed along the A-P axis in *Ciona.* From these neural progenitors expressing different combinations, or codes, of these homeodomain proteins, the 5 distinct neuronal subtypes of the *Ciona* MG are born. We sought to characterize the expression patterns of homologous genes in the developing MG of *M. occidentalis.* We were able to identify these readily by performing BLAST against our transcriptome assembly, which revealed clear orthologs for the following genes: *Mnx, Vsx, Islet, Nk6, Pax3/7*, and *Dmbx.* An additional gene, *Lhx3/4.a* had already been identified and characterized in our previous study (Stolfi et al., 2014).

### Motor neuron and interneuron identities

In *Ciona*, 4 of the 5 core MG neuron subtypes are marked by either *Mnx* or *Vsx* (Imai et al., 2009; Stolfi and Levine, 2011). These are orthologs of conserved transcription factors that specify motor neurons (HB9, MNR2, etc.) or interneurons (Chox10, CHX10, Ceh-10, etc.), respectively. Two-color ISH revealed a neatly alternating *Vsx-Mnx-Vsx-Mnx* pattern on one side of the developing MG of *M. occidentalis*, around 8.5 hours post-fertilization (hpf)(Fig 4A). This alternating pattern closely mirrors the alternation of motor neuron (MN) and interneuron (IN) subtypes in the *Ciona* MG: IN1-MN1-IN2-MN2. In the *Ciona* MG, the latter three are the first MG neurons to differentiate, corresponding to cells A11.118 (MN1), A11.117 (IN2), and A10.64 (MN2), which are consecutively arrayed with no intervening cells between them. This appears to be the case in *M. occidentalis* also. However, in *Ciona* there are additional non-neuronal cells intercalated between IN1 and MN1, as a result of continued proliferation in the anterior MG that ultimately gives rise to IN1 (see Fig 2B). In *Ciona*, *Vsx* expression is restricted to post-mitotic interneurons, and is not observed in A11.119, the grandmother cell of IN1 (Stolfi and Levine, 2011). In *M. occidentalis*, the lack of any discernible gap between the anterior-most *Vsx+* and *Mnx+* cells at first suggests that *Vsx* transcription starts in the A11.119 progenitor cell itself, a clear example of transcriptional “priming” of cell fate (Razy-Krajka et al., 2014). Indeed, we observed *Vsx* expression in two anterior cells when observed at slightly later stages, suggesting that A11.119 divided after the onset of *Vsx* activation (Fig 4B). Together with evidence for earlier *Dmbx* expression in the A11.120 progenitor cell (see below), these data suggest that the expression of certain MG neuron subtype-specific transcription factors is already primed in *M. occidentalis* MG progenitors. This heterochronic shift may be related to the ~10% faster development of *Molgula* relative to *Ciona* (Stolfi et al., 2014).

**Figure 4.**
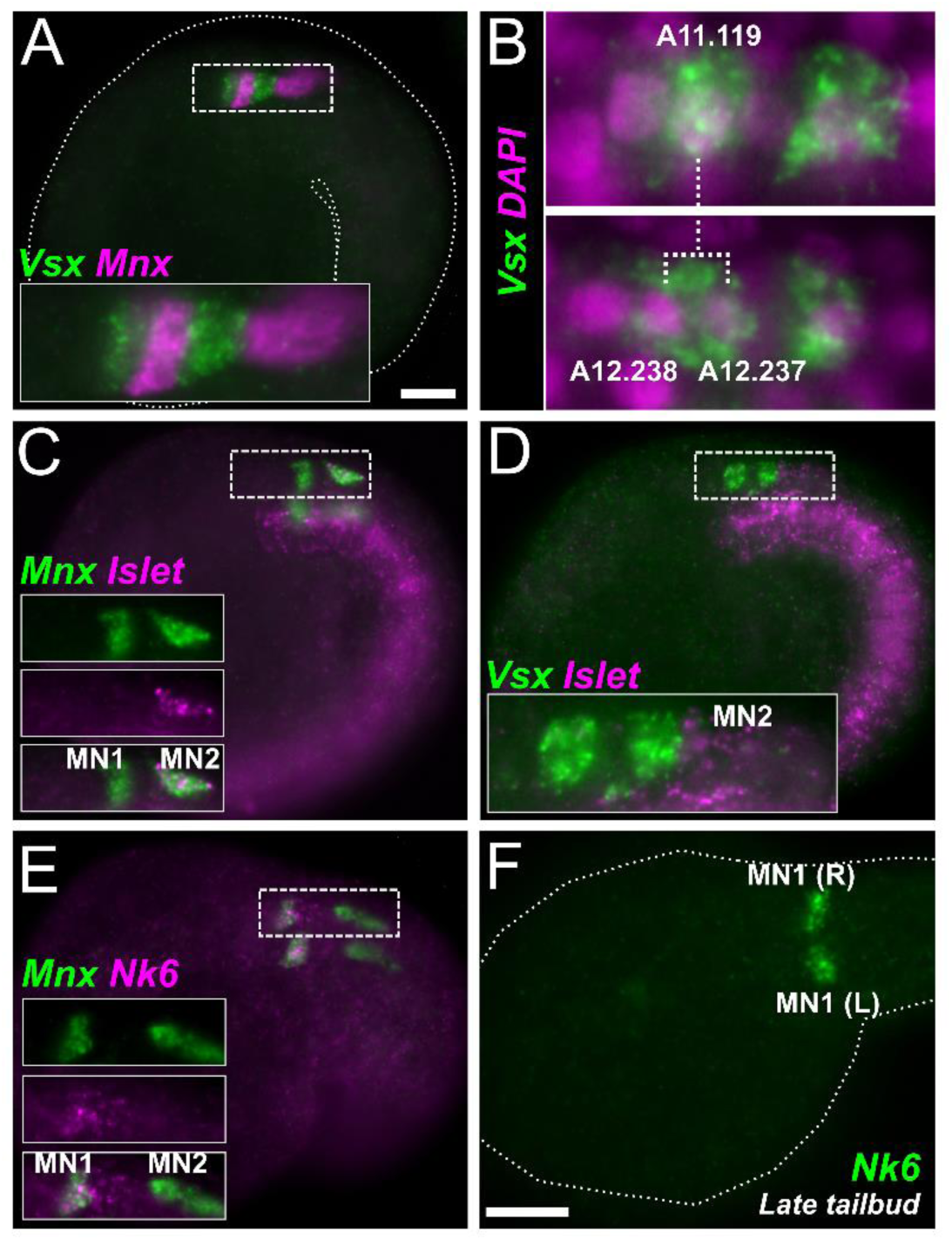
Motor neurons and interneurons in the MG. **A)** Lateral view of mid-tailbud stage *Molgula occidentalis* embryo (dotted outline). Two-color fluorescent *in situ* hybridization reveals expression of interneuron marker *Vsx* (green) and motor neuron marker *Mnx* (magenta) in alternating cells in the MG (only one side shown). Inset is a magnified view of dashed box. **B)** *In situ* hybridization for *Vsx* (green) in embryos at two successive stages. *Vsx* is initially activated in A11.119 (top panel), which divides and gives rise to daughter cells A12.238 and A12.237, both of which continue to express *Vsx* at this stage. These data confirm that *Vsx* is activated in the A11.119 cell in *M. occidentalis*, which is not the case in *Ciona* (see text for details). Nuclei are counterstained with DAPI (magenta). **C)** Two-color fluorescent *in situ* hybridization for *Mnx* (green) and MN2 marker *Islet* (magenta), revealing identities of MN1 (*Mnx+/Islet-)* and MN2 (*Mnx+/Islet+)* cells. Boxed area magnified in insets. **D)** Two-color fluorescent *in situ* hybridization for *Vsx* (green) and *Islet* (magenta). Only one side of embryo is in focus. **E)** Two-color fluorescent *in situ* hybridization for *Mnx* (green) and MN1 marker *Nk6* (magenta), confirming identities of MN1 (*Mnx+/Nk6+)* and MN2 (*Mnx+/Nk6-).* **F)** *Nk6 in situ* hybridization (green) in late tailbud embryo, showing sharpened expression in MN1 cells on both sides of the MG (R = right side, L = left side). All scale bars = 25 µm.

### *Islet* expression reveals MN2 (A10.64 cell)

From the *Mnx* ISH, MN2 appeared to be the posterior-most cell of the core MG. The identity of this cell was confirmed by two-color ISH with *Islet*, a marker of MN2 fate in *Ciona* (Fig 4C,D). In *Ciona*, MN2 is the only core MG neuron that is not derived from the A9.30 lineage. Long identified as the A10.57 cell derived from the A9.29 lineage, MN2 was recently revealed in fact to be the A10.64 cell of the A9.32 lineage instead and ultimately derived from the A8.16 neuromesodermal lineage that also gives rise to the secondary tail muscles of the larva (Navarrete and Levine, 2016). Despite its origin from further posterior in the embryo, MN2 becomes associated with the MG thanks to a dramatic, anterior migration along the outside of the neural tube, leap-frogging over the entire A9.29 lineage (Navarrete and Levine, 2016). The migrating MN2 is elongated along the A-P axis due to the extension of a leading edge that ultimately contacts the A9.30 lineage. Upon contacting the A9.30 lineage, MN2 extends its axon posteriorly, retaining an elongated cell body and forming *en passant* synapses with the dorsal band of muscle cells down the length of the tail (Imai and Meinertzhagen, 2007). In *M. occidentalis*, the elongated morphology of MN2 was readily apparent by ISH, in which fluorescent signal fills most of the cell bodies to reveal their shapes. Therefore we conclude that, in *M. occidentalis*, the specification and morphogenesis of MN2 is highly conserved.

### *Nk6* expression is refined and restricted to MN1

While MN2 controls tail contractions in a graded manner (Nishino et al., 2011), the other major MN in *Ciona* is MN1, the A11.118 cell. This neuron was shown to form large, leaf-like (“frondose”) motor endplates at the base of the tail (Stolfi and Levine, 2011). While MN2s are proposed to exert graded motor control during maneuvering, the all-or-none flexions of the tail that drive swimming are thought to be triggered instead by MN1s. This cell is characterized by *Mnx* expression without co-expression of *Islet.* In *Ciona*, MN1 is also marked by late, sustained expression of *Nk6* close to hatching, even though this gene is expressed broadly throughout the posterior A9.30 lineage earlier in development (Stolfi et al., 2011). In *M. occidentalis*, we found this also to be true. *Nk6* expression was seen in both MN1 and IN2 at the mid tailbud stage (Fig 4D), but later was seen only in a single pair of cells in the embryo, presumably MN1s (Fig 4E). This further confirms the highly conserved nature of the posterior MG, and the specification of MN1 by sustained *Nk6* expression.

### Subfunctionalization of *Lhx3/4* paralogs

In *Ciona* and the stolidobranch tunicate *Halocynthia roretzi*, a single *Lhx3/4* is transcribed as two alternative isoforms, transcript variants 1 and 2, originating from alternate promoters (Fig 5A)(Christiaen et al., 2009a; Kobayashi et al., 2010). In both *Ciona* and *Halocynthia*, transcript variant 1 is expressed in MG precursors, while transcript variant 2 is expressed in early vegetal pole cells and is required for endoderm and cardiopharyngeal mesoderm specification (Christiaen et al., 2009a; Takatori et al., 2010). We previously found that in all three *Molgula* genomes sequenced to date, the ancestral tunicate *Lhx3/4* gene was duplicated, resulting in two paralogs, termed *Lhx3/4.a* and *Lhx3/4.b* (Fig 5A)(Stolfi et al., 2014). In *M. occidentalis*, their expression patterns and putative functions appear to mirror those of the two different transcript variants identified in *Ciona/Halocynthia: Lhx3/4.a* is expressed in the MG while *Lhx3/4.b* is expressed in the early vegetal pole cells and is involved in cardiopharyngeal mesoderm specification (Stolfi et al., 2014). *Lhx3/4.a* is the more conservative paralog, retaining an ancestral C-terminal peptide motif, which has been lost in *Lhx3/4.b* (Stolfi et al., 2014). Given that *Halocynthia* is more closely related to *Molgula* than to *Ciona* (Tsagkogeorga et al., 2009), this suggests a *Molgula-*specific duplication of *Lhx3/4*, followed by subfunctionalization through partitioning of the expression domains of the ancestral transcript variants in addition to protein-coding changes.

**Figure 5.**
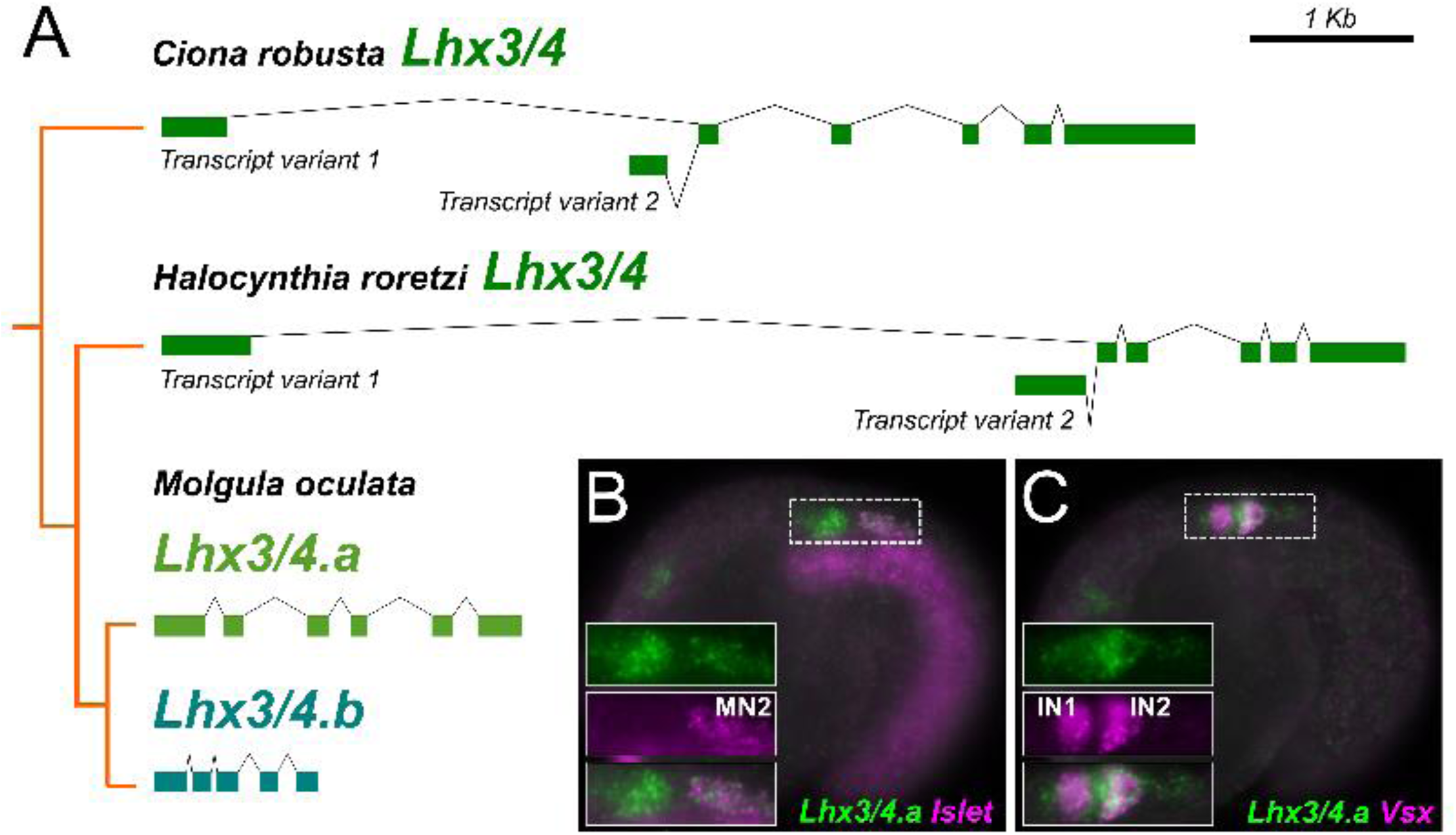
*Molgula-*specific duplication and sub-functionalization of *Lhx3/4*. **A)** Diagram depicting *Lhx3/4* genes in the genomes of *Ciona robusta, Halocynthia roretzi*, and *Molgula oculata*, with their phylogenetic relationships indicated by the orange tree. All diagrams are at the same scale (scale bar is at top right). Exons indicated as colored bars, intron are depicted by crooked lines. Both *Ciona* and *Halocynthia* have a single *Lhx3/4* gene that is transcribed from two distinct promoters to give rise to two transcript variants. In both species, variant 2 is involved in vegetal pole patterning in the early embryo, while variant 1 is involved in motor ganglion patterning. In all three *Molgula* species sequenced, *Lhx3/4* has been duplicated, giving rise to paralogs *Lhx3/4.a* and *Lhx3/4.b.* Loci from *M. oculata* are shown here (instead of *M. occidentalis*) because its genome is the least fragmented *Molgula* genome. *M. occidentalis Lhx3/4* genes are similar in structure to their *M. oculata* orthologs, but assembly errors near the 3’ end of *M. occidentalis Lhx3/4.b* (not shown) prevented us from using sequences from this species in our diagram. *Ciona* and *Molgula* data are from Christiaen et al. 2009a and Kobayashi et al. 2010. **B)** Two-color fluorescent *in situ* hybridization for *Lhx3/4.a* (green) and *Islet* (magenta) showing co-expression in MN2. **C)** Two-color fluorescent *in situ* hybridization for *Lhx3/4.a* (green) and *Vsx* (magenta) showing strong co-expression in IN2, and diffuse *Lhx3/4.a* expression throughout the MG.

Here we looked more closely at the expression of *Lhx3/4.a* in the developing MG of *M. occidentalis*. In *Ciona, Lhx3/4* is initially expressed in the posterior MG (MN1, IN2, and more weakly in MN2), and later in IN1 (Ikuta and Saiga, 2007a; Imai et al., 2009). In *Halocynthia, Lhx3/4* was also detected earlier in development, in the A9.30 blastomere (Katsuyama et al., 2005). In *M. occidentalis*, we found *Lhx3/4.a* expressed throughout the entire MG at the mid-tailbud stage, though more strongly in MN1 and IN2 (Fig 5B,C). Higher expression in these cells is shared with *Ciona* and *Halocynthia*, and the weaker expression in the anterior MG may reflect fading expression from earlier stages as in *Halocynthia.* Based on these results, we propose that *Lhx3/4.a* has, for the most part, conserved the MG-specific expression pattern and putative function of the ancestral *Lhx3/4* gene, more specifically those of its transcript variant 1.

### *Pax3/7* and *Dmbx* delineate the descending decussating neurons (ddNs)

The anterior-most neuron in the core MG of *Ciona* is the descending decussating neuron (ddN), corresponding to the A12.239 cell. The recently mapped *Ciona intestinalis* connectome revealed that this neurons are likely homologous to giant reticulospinal Mauthner cells (M-cells) of vertebrates and situated in a neural circuit that is topologically very similar to M-cell escape circuits (Ryan et al., 2017). In *Ciona*, the ddN is specified by a gene regulatory network involving the transcription factors Pax3/7 and Dmbx (Stolfi et al., 2011). To summarize, Pax3/7 is sufficient and necessary for the specification of the A11.120 progenitor, the mother cell of the ddN, and directly activates *Dmbx* expression in the posterior daughter cell of A11.120, which becomes the ddN (A12.239). *Dmbx* in turn encodes a repressor that mediates an FGF-regulated switch for differentiation (Stolfi et al., 2011). Upon differentiating, ddNs project their nascent axons across the midline, perpendicular to the A-P axis. Thus, as their name implies, they are the only core MG neurons that decussate.

We therefore sought to characterize the expression patterns of *Pax3/7* and *Dmbx* in *M. occidentalis.* ISH revealed *Pax3/7* and *Dmbx* expression in a single cell in the MG, just anterior to the *Vsx*-expressing cell identified as A11.119 (Fig 6A,B). These data indicate that *Pax3/7* and *Dmbx* are co-expressed in A11.120, the mother cell of the ddN. While *Pax3/7* expression in A11.120 is shared between *Molgula* and *Ciona, Dmbx* expression in *M. occidentalis* appears to be shifted earlier relative to mitotic exit of the neuron. In *Ciona*, transcription of *Dmbx* is not detected in the A11.120 cell but is activated later, specifically in the ddN, by combinatorial action of Pax3/7 (inherited from the mother cell), and Neurogenin (Stolfi et al., 2011).

**Figure 6.**
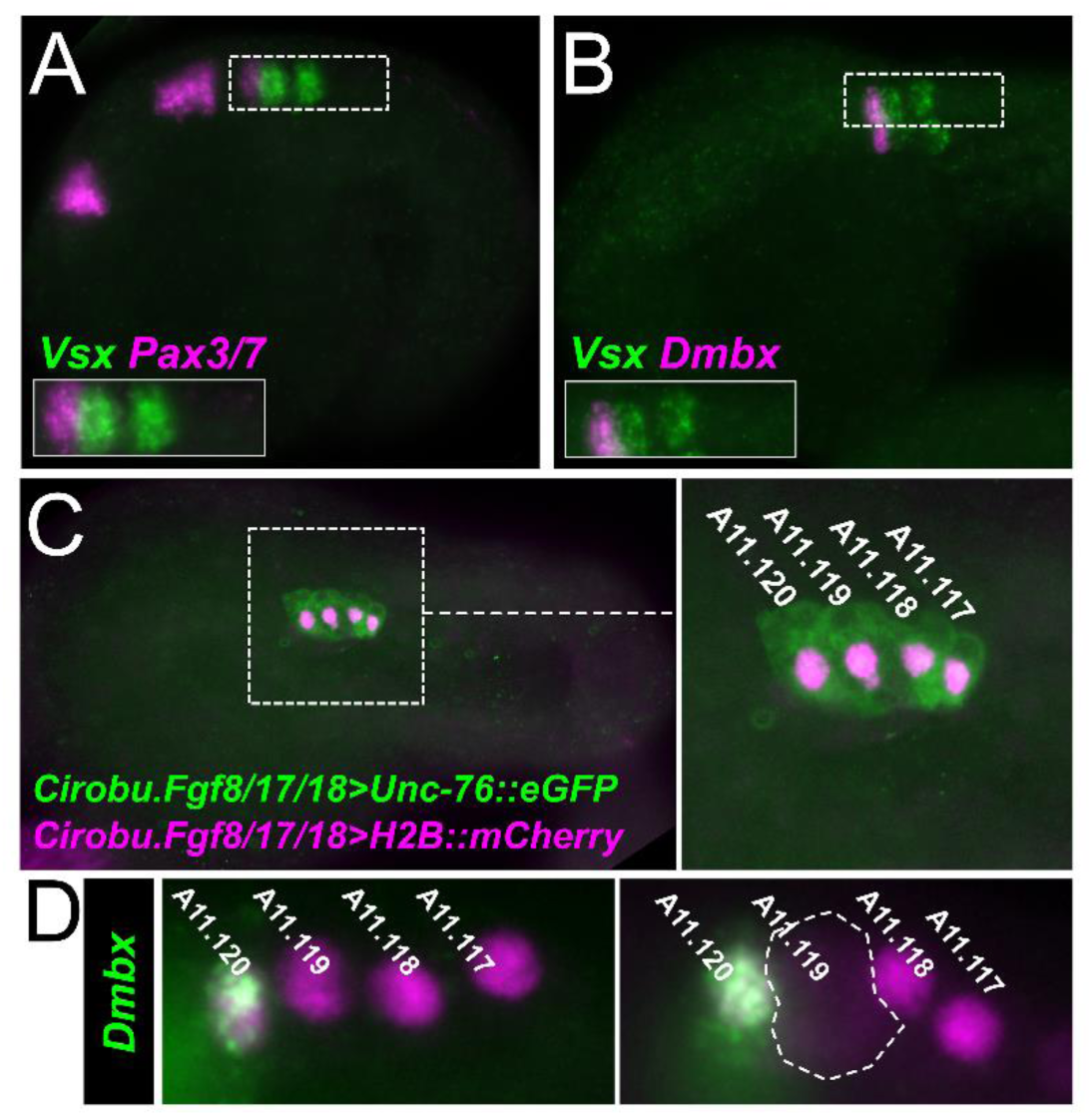
*Pax3/7* and *Dmbx* activation in A11.120. **A)** Two-color fluorescent *in situ* hybridization for *Vsx* (green) and *Pax3/7* (magenta) showing expression of *Pax3/7* anterior to *Vsx* expression in A11.119 cell. **B)** Two-color fluorescent *in situ* hybridization for *Vsx* (green) and *Dmbx* (magenta) also showing expression of *Dmbx* anterior to *Vsx* expression in A11.119 cell. **C)** *M. occidentalis* early tailbud embryo electroporated with *Ciona robusta* (*Cirobu) Fgf8/17/18* reporter plasmids, labeling the A9.30 lineage on one side, with H2B::mCherry-labeled nuclei in magenta and Unc-76::eGFP-labeled cell bodies in green. Inset at right showing higher magnification of boxed area, showing all four descendants of A9.30, arrayed from anterior to posterior. **D)** *In situ* hybridization of *Dmbx* coupled to immunostaining of Beta-galactosidase (magenta nuclei) driven by *Cirobu.Fgf8/17/18>nls::lacZ* reporter plasmid. *Dmbx* is strongly activated in the A11.120 cell, the mother cell of the descending decussating neuron (ddN). In *Ciona*, *Dmbx* is detected only in the differentiating ddN, not in A11.120. Right panel shows A11.119 entering mitosis indicated by nuclear envelope breakdown and diffusion on nuclearly-localized Beta-galactosidase (dashed outline). A11.119 appears to divide before the other cells in the MG at this stage, which is also the case in *Ciona.* This further confirms the precocious activation of *Vsx* in this cell relative to *Ciona* (see Figure 4).

To confirm that *Dmbx* is expressed in the A11.120 cells of *M. occidentalis*, we sought to determine its expression pattern at cellular resolution. We turned to electroporation to transfect embryos with a reporter construct that could selectively label the A9.30 lineage and allow us to combine ISH with immunofluorescence. To this end, we used the *Fgf8/17/18* promoter from *C.* robusta (Imai et al., 2009) to drive expression of Histone 2B::mCherry (*Cirobu.Fgf8/17/18>H2B::mCherry*) in the A9.30 lineage of electroporated *M. occidentalis* embryos. Activity of this reporter plasmid was relatively weak and infrequent in *M. occidentalis* embryos, but managed to label the 4 cells of the developing MG of *M. occidentalis*, equivalent to the cells A11.120, A11.119, A11.118, and A11.117 (from anterior to posterior)(Fig 6C), and ISH + immunofluorescence revealed *Dmbx* expression in A11.120 at this stage (Fig 6D). Taken together, these data indicate a *M. occidentalis-*specific priming of *Dmbx* transcription in the progenitor of the ddN, similar to the priming of *Vsx* in the progenitor of IN1 in this species as well (see above).

### Conserved morphogenesis and axon projection of ddNs in *M. occidentalis*

Given the unique morphology and axon trajectory of the ddNs in *Ciona* and their likely homology to M-cells in a conserved, pan-chordate escape network (Ryan et al., 2017), we asked whether their morphogenesis is conserved in *M. occidentalis.* To label differentiated ddNs, we electroporated *M. occidentalis* embryos with a *Dmbx* reporter plasmid containing ~1.4 Kb of genomic DNA sequence 5’ to the ATG corresponding to the predicted start codon of the *Dmbx* mRNA from the related species *M. oculata* (**Supplemental File 1**), fused to the *Unc-76::eGFP* fluorescent reporter gene to label axons uniformly (Dynes and Ngai, 1998). This construct was sufficient to label the ddNs of *M. occidentalis* (Fig 7A,B). Unc-76::eGFP expression in the ddNs revealed a polarization along the medial-lateral axis, perpendicular to the A-P, resulting in their unique axon trajectory initially straight across the midline, then abruptly turning 90° to descend along the outside of the neural tube towards the tail. This closely mirrors the processes of axonogenesis and axon guidance that result in the conserved axon trajectory of the ddNs in *Ciona* (Stolfi and Levine, 2011). This suggests that ddN morphogenesis is an ancient and highly conserved process in tunicates, which also shares features with M-cell morphogenesis in vertebrates (Kimmel and Model, 1978).

**Figure 7.**
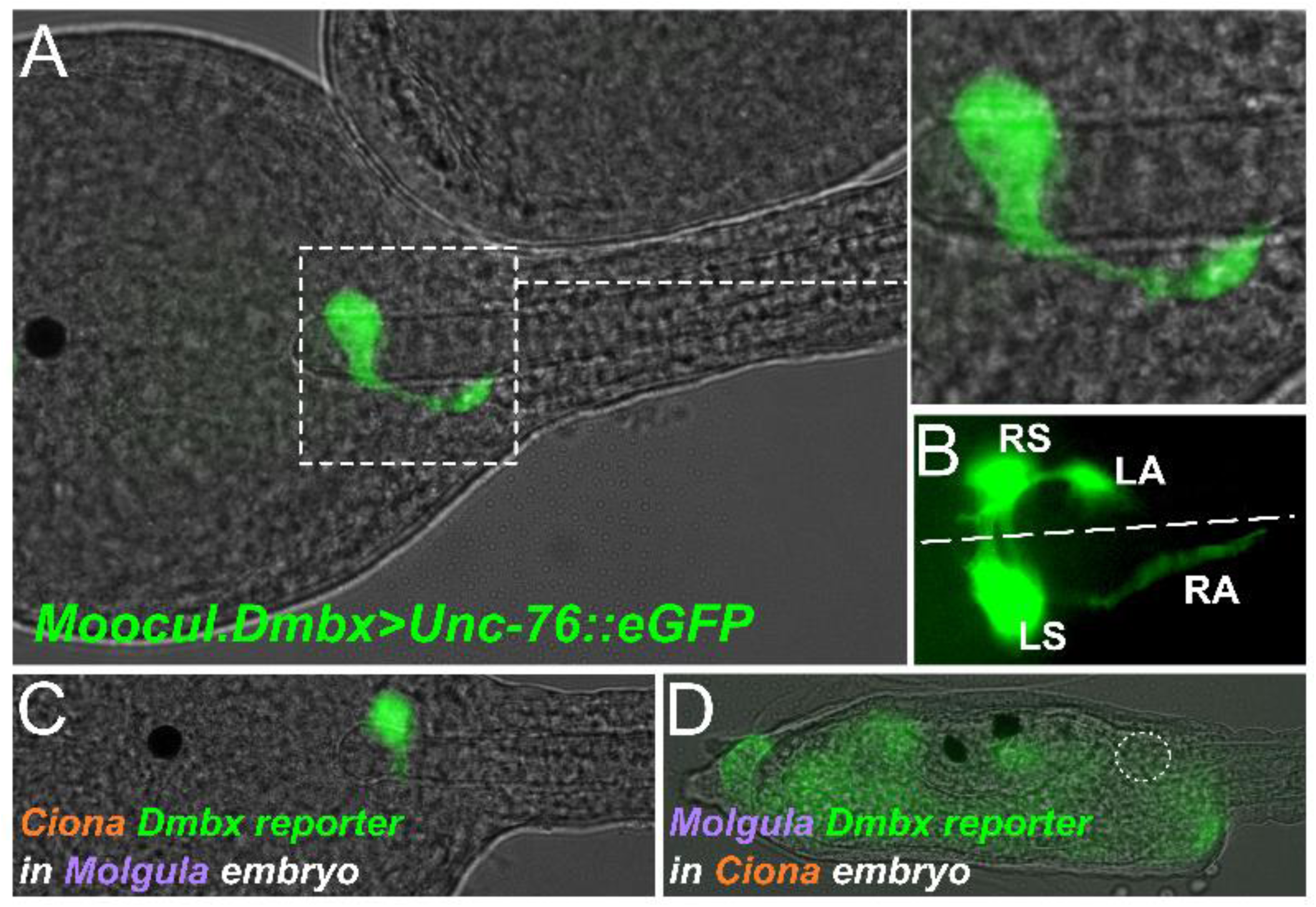
*Dmbx* reporter assays reveal conserved morphogenesis of the ddN and asymmetric unintelligibility of *cis*-regulatory sequences. **A)** Dorsal view of a *Molgula occidentalis* embryo electroporated with a *Molgula oculata (Moocul) Dmbx>Unc-76::eGFP* reporter plasmid, specifically labeling (in green) a developing descending decussating neuron (ddN) on one side of the embryo. Inset at top right shows magnified view of boxed area, showing the conserved axon trajectory of the ddN, which is initially perpendicular to the anterior-posterior axis and then turns 90 degrees to continue posteriorly towards the tail. **B)** Another embryo electroporated with *Moocul.Dmbx>Unc-76::eGFP*, this time labeling both right and left ddNs. Both axons cross the midline (dashed line) to continue down towards the tail on the other side. RS: right soma, RA: right axon, LS: left soma, LA: left axon. **C)** *M. occidentalis* embryo electroporated with *Ciona robusta Dmbx>Unc-76::Venus* reporter, which is activated precisely and robustly in the ddN (green). **D)** *Ciona robusta* embryo electroporated with *Molgula oculata Dmbx>Unc-76::eGFP* reporter, which is not active in this species’ ddN (approximate location indicated by dashed circle). Faint, leaky expression (green) can be seen in other unrelated tissues.

### Cross-species *Dmbx* reporter assays reveal asymmetric developmental system drift (DSD)

In our comparison of cardiopharyngeal development between *Ciona* and *Molgula*, we discovered pervasive, acute developmental system drift (DSD)(True and Haag, 2001) that has resulted in cross-species incompatibility of orthologous *cis-*regulatory DNAs that regulate identical gene expression patterns in the tunicate cardiopharyngeal mesoderm (Stolfi et al., 2014). Here we tested whether DSD might also underlie the highly conserved gene expression patterns seen in the developing MG. We found that the previously identified *C. robusta Dmbx* driver (Stolfi and Levine, 2011) was sufficient to activate reporter gene expression in *M. occidentalis* ddNs (Fig 7C). However, the *M. oculata Dmbx* reporter plasmid, used to label *M. occidentalis* ddNs, was non-functional in *C. robusta* embryos (Fig 7D). Thus, there is an asymmetry in the intelligibility of *cis-*regulatory logic for *Dmbx* expression*; Ciona* logic works in *Molgula*, but not vice-versa.

One possible explanation for this asymmetry is that, in *Molgula, Dmbx* is initially activated in the A11.120 cell, the mother cell of the ddN (see Fig 6D). In *Ciona*, *Dmbx* is only detected later, in the post-mitotic ddN (Stolfi and Levine, 2011), and is activated by a combination of Neurogenin and Pax3/7, and depends critically on the presence of a Pax3/7 binding site located in a *cis*-regulatory element that controls *Dmbx* transcription in the ddN (Stolfi et al., 2011). Therefore, it is possible that an alternate logic drives the earlier activation of *Dmbx* that we observe in *Molgula*, considering that *Pax3/7* and *Dmbx* are transcribed simultaneously in *M. occidentalis* MG, not sequentially as in *Ciona*. The required factors for this earlier, Pax3/7-independent activation of *Dmbx* in *M. occidentalis* might therefore be absent from the *Ciona* MG, resulting in lack of activation of the *Molgula* reporters in *Ciona.* In contrast, we show that both *Pax3/7* (see Fig 6A) and *Neurogenin* (see Fig 3B) expression patterns are conserved in *M. occidentalis*, which would allow for proper activation of the *Ciona Dmbx* reporter plasmid in this species.

## Conclusions

Here we have documented the deep conservation of regulatory gene expression patterns that establish the precise configuration of neuronal subtypes in the MG of the tunicate larva. Our data suggest that the circuitry of the MG, as revealed by the recently completed *C. intestinalis* connectome [connectome elife], may be as rigidly conserved as the embryonic development of the larva itself. We did not observe any indication that certain MG neuron subtypes identified in *Ciona* are missing in *M. occidentalis*, or that additional subtypes (not observed in *Ciona*) are specified in *M. occidentalis.* Not only does there appear to be a 1-to-1 correspondence of core MG neurons between *Ciona* and *M. occidentalis*, their position in the MG is not altered. The only differences observed are precocious transcription of certain markers in *M. occidentalis* (e.g. *Vsx, Dmbx*), which may be related to the faster developmental rate of this species relative to *Ciona* spp. These observations suggest that the MG of the solitary tunicate larva has changed very little in nearly 400 million years, the estimated time of divergence between *Ciona* and *Molgula* (Delsuc et al., 2018). As such, the tunicate larval MG likely represents a minimal but ancient and exquisitely adapted central pattern generator for swimming behavior.

Our finding of acute, asymmetric DSD of *Dmbx* regulation hints at a possible cause for divergence in the regulation of seemingly identical gene expression patterns. In this case, both *C. robusta* and *M. occidentalis* specify a single pair of *Dmbx+* ddNs. However, precocious transcriptional priming of *Dmbx* in the ddN progenitor (A11.120) in *M. occidentalis* may be the result of an alternate regulatory mechanism that does not operate in homologous cells of *Ciona.* Thus, while the spatial pattern of ddN specification is highly conserved, likely constrained by the invariance of the MG and its developmental lineages, there have been a changes to the temporal dynamics of this process. Although the developmental timing of the ddN is identical between *Molgula* and *Ciona* in terms of number of mitotic generations, *Molgula* development is accelerated on an absolute time scale (e.g. hours post-fertilization). This accelerated developmental rate on an absolute timescale may have required very different regulatory strategies simply to maintain the same output (i.e. specification of the A12.239 pair of cells as the ddNs). It will be interesting to investigate in the future whether the root of the many *cis*/*trans* incompatibilities observed between these geometrically identical embryos lies mostly in their different absolute developmental rates.

The remarkable conservation of embryonic cell lineages and geometries between distantly related tunicates has been recently proposed to also underlie the extreme divergence of tunicate genomes (Guignard et al., 2018). Because gene expression and cell fates are induced by very precise and invariant cell-cell contact events in tunicate embryos, the gene regulatory networks involved in their regulation have been allowed to drift, freed from the constraints imposed by the mechanisms required by larger, more complex and variable embryos. In other words, precision and robustness of gene expression in tunicates arise from the precision and robustness of the embryo itself and not its gene networks. In larger animals (e.g. vertebrates), this precision and robustness can only come from more precise and robust gene regulatory networks instead. Additionally, while the invariant geometry of the tunicate embryo may have served to release the genome from these constraints, it may have imposed a different set of evolutionary constraints. Because of their invariance, tunicate embryos are unable to compensate for errors in cell fate specification and likely experienced strong selective pressure to maintain precise embryo geometries even as some evolved to develop more rapidly on an absolute timescale, like *M. occidentalis*. It is perhaps due to this evolutionary ratchet that *Oikopleura dioica*, the tunicate with the fastest rate of development and most geometrically constrained embryo, also has the most fragmented, highly derived genome as well (Denoeud et al., 2010; Edvardsen et al., 2004; Fujii et al., 2008; Nishida, 2008; Seo et al., 2004; Seo et al., 2001). Future comparative studies among the tunicates promise to shed further light on the fascinating interplay between embryonic form and genetic architecture in evolution.

## Acknowledgments

We thank Rodoniki Athanasiadou and David Gresham for helping us with the RNAseq. We are also grateful for the constant support from the MoEvoDevo collaborative network: Billie Swalla, C. Titus Brown, Lionel Christiaen, Filomena Ristoratore, and Claudia Racioppi. Work in the A.S. laboratory is funded by NIH award R00 HD084814. The authors have no competing interests to declare.

## References

1. Satoh N: Developmental genomics of ascidians: John Wiley & Sons; 2013.

2. Nishino A: Morphology and Physiology of the Ascidian Nervous Systems and the Effectors. In: Transgenic Ascidians. Springer; 2018: 179–196.

3. White JG, Southgate E, Thomson JN, Brenner S: The structure of the nervous system of the nematode Caenorhabditis elegans. Philos Trans R Soc Lond B Biol Sci 1986, 314(1165):1–340.

4. Ryan K, Lu Z, Meinertzhagen IA: The CNS connectome of a tadpole larva of Ciona intestinalis (L.) highlights sidedness in the brain of a chordate sibling. Elife 2016, 5.

5. Ryan K, Lu Z, Meinertzhagen IA: The peripheral nervous system of the ascidian tadpole larva: Types of neurons and their synaptic networks. Journal of Comparative Neurology 2018, 526(4):583–608.

6. Ryan K, Lu Z, Meinertzhagen IA: Circuit homology between decussating pathways in the Ciona larval CNS and the vertebrate startle-response pathway. Current Biology 2017, 27(5):721–728.

7. Nishino A, Okamura Y, Piscopo S, Brown ER: A glycine receptor is involved in the organization of swimming movements in an invertebrate chordate. BMC neuroscience 2010, 11(1):6.

8. Wada H, Saiga H, Satoh N, Holland PW: Tripartite organization of the ancestral chordate brain and the antiquity of placodes: insights from ascidian Pax-2/5/8, Hox and Otx genes. Development 1998, 125(6):1113–1122.

9. Imai KS, Stolfi A, Levine M, Satou Y: Gene regulatory networks underlying the compartmentalization of the Ciona central nervous system. Development 2009, 136(2):285–293.

10. Stolfi A, Levine M: Neuronal subtype specification in the spinal cord of a protovertebrate. Development 2011, 138(5):995–1004.

11. Stolfi A, Wagner E, Taliaferro JM, Chou S, Levine M: Neural tube patterning by Ephrin, FGF and Notch signaling relays. Development 2011, 138(24):5429–5439.

12. Hudson C, Ba M, Rouvière C, Yasuo H: Divergent mechanisms specify chordate motoneurons: evidence from ascidians. Development 2011, 138(8):1643–1652.

13. Tsagkogeorga G, Cahais V, Galtier N: The population genomics of a fast evolver: high levels of diversity, functional constraint, and molecular adaptation in the tunicate Ciona intestinalis. Genome Biology and Evolution 2012, 4(8):852–861.

14. Tsagkogeorga G, Turon X, Galtier N, Douzery EJ, Delsuc F: Accelerated evolutionary rate of housekeeping genes in tunicates. Journal of molecular evolution 2010, 71(2):153–167.

15. Delsuc F, Philippe H, Tsagkogeorga G, Simion P, Tilak M-K, Turon X, López-Legentil S, Piette J, Lemaire P, Douzery EJ: A phylogenomic framework and timescale for comparative studies of tunicates. BMC biology 2018, 16(1):39.

16. Kocot KM, Tassia MG, Halanych KM, Swalla BJ: Phylogenomics offers resolution of major tunicate relationships. Molecular phylogenetics and evolution 2018.

17. Stolfi A, Lowe EK, Racioppi C, Ristoratore F, Brown CT, Swalla BJ, Christiaen L: Divergent mechanisms regulate conserved cardiopharyngeal development and gene expression in distantly related ascidians. Elife 2014, 3.

18. True JR, Haag ES: Developmental system drift and flexibility in evolutionary trajectories. Evolution & development 2001, 3(2):109–119.

19. Guignard L, Fiuza U-M, Leggio B, Faure E, Laussu J, Hufnagel L, Malandain G, Godin C, Lemaire P: Contact-dependent cell communications drive morphological invariance during ascidian embryogenesis. bioRxiv 2018: 238741.

20. Brozovic M, Martin C, Dantec C, Dauga D, Mendez M, Simion P, Percher M, Laporte B, Scornavacca C, Di Gregorio A: ANISEED 2015: a digital framework for the comparative developmental biology of ascidians. Nucleic acids research 2015, 44(D1):D808–D818.

21. Brozovic M, Dantec C, Dardaillon J, Dauga D, Faure E, Gineste M, Louis A, Naville M, Nitta KR, Piette J: ANISEED 2017: extending the integrated ascidian database to the exploration and evolutionary comparison of genome-scale datasets. Nucleic acids research 2017, 46(D1):D718–D725.

22. Crusoe MR, Alameldin HF, Awad S, Boucher E, Caldwell A, Cartwright R, Charbonneau A, Constantinides B, Edvenson G, Fay S: The khmer software package: enabling efficient nucleotide sequence analysis. F1000Research 2015, 4.

23. Schulz MH, Zerbino DR, Vingron M, Birney E: Oases: robust de novo RNA-seq assembly across the dynamic range of expression levels. Bioinformatics 2012, 28(8):1086–1092.

24. Pryszcz LP, Gabaldón T: Redundans: an assembly pipeline for highly heterozygous genomes. Nucleic acids research 2016, 44(12):e113–e113.

25. Hunt M, Kikuchi T, Sanders M, Newbold C, Berriman M, Otto TD: REAPR: a universal tool for genome assembly evaluation. Genome biology 2013, 14(5):R47.

26. Neymotin B, Athanasiadou R, Gresham D: Determination of in vivo RNA kinetics using RATE-seq. Rna 2014, 20(10):1645–1652.

27. Parkhomchuk D, Borodina T, Amstislavskiy V, Banaru M, Hallen L, Krobitsch S, Lehrach H, Soldatov A: Transcriptome analysis by strand-specific sequencing of complementary DNA. Nucleic acids research 2009, 37(18):e123–e123.

28. Bolger AM, Lohse M, Usadel B: Trimmomatic: a flexible trimmer for Illumina sequence data. Bioinformatics 2014, 30(15):2114–2120.

29. Kim D, Langmead B, Salzberg SL: HISAT: a fast spliced aligner with low memory requirements. Nature methods 2015, 12(4):357.

30. Li H, Handsaker B, Wysoker A, Fennell T, Ruan J, Homer N, Marth G, Abecasis G, Durbin R: The sequence alignment/map format and SAMtools. Bioinformatics 2009, 25(16):2078–2079.

31. Pertea M, Pertea GM, Antonescu CM, Chang T-C, Mendell JT, Salzberg SL: StringTie enables improved reconstruction of a transcriptome from RNA-seq reads. Nature biotechnology 2015, 33(3):290.

32. Trapnell C, Roberts A, Goff L, Pertea G, Kim D, Kelley DR, Pimentel H, Salzberg SL, Rinn JL, Pachter L: Differential gene and transcript expression analysis of RNA-seq experiments with TopHat and Cufflinks. Nature protocols 2012, 7(3):562.

33. Simão FA, Waterhouse RM, Ioannidis P, Kriventseva EV, Zdobnov EM: BUSCO: assessing genome assembly and annotation completeness with single-copy orthologs. Bioinformatics 2015, 31(19):3210–3212.

34. Christiaen L, Wagner E, Shi W, Levine M: Isolation of sea squirt (Ciona) gametes, fertilization, dechorionation, and development. Cold Spring Harbor Protocols 2009, 2009(12):pdb. prot5344.

35. Ikuta T, Saiga H: Dynamic change in the expression of developmental genes in the ascidian central nervous system: revisit to the tripartite model and the origin of the midbrain-hindbrain boundary region. Developmental biology 2007, 312(2):631–643.

36. Cole AG, Meinertzhagen IA: The central nervous system of the ascidian larva: mitotic history of cells forming the neural tube in late embryonic Ciona intestinalis. Developmental biology 2004, 271(2):239–262.

37. Nicol D, Meinertzhagen I: Development of the central nervous system of the larva of the ascidian, Ciona intestinalis L: I. The early lineages of the neural plate. Developmental biology 1988, 130(2):721–736.

38. Nicol D, Meinertzhagen I: Development of the central nervous system of the larva of the ascidian, Ciona intestinalis L: II. Neural plate morphogenesis and cell lineages during neurulation. Developmental biology 1988, 130(2):737–766.

39. Navarrete IA, Levine M: Nodal and FGF coordinate ascidian neural tube morphogenesis. Development 2016, 143(24):4665–4675.

40. Nishitsuji K, Horie T, Ichinose A, Sasakura Y, Yasuo H, Kusakabe TG: Cell lineage and cis‐regulation for a unique GABAergic/glycinergic neuron type in the larval nerve cord of the ascidian Ciona intestinalis. Development, growth & differentiation 2012, 54(2):177–186.

41. Satou Y, Takatori N, Yamada L, Mochizuki Y, Hamaguchi M, Ishikawa H, Chiba S, Imai K, Kano S, Murakami SD et al: Gene expression profiles in *Ciona intestinalis* tailbud embryos. Development 2001, 128(15):2893–2904.

42. Mazet F, Hutt JA, Milloz J, Millard J, Graham A, Shimeld SM: Molecular evidence from Ciona intestinalis for the evolutionary origin of vertebrate sensory placodes. Developmental biology 2005, 282(2):494–508.

43. Takamura K, Egawa T, Ohnishi S, Okada T, Fukuoka T: Developmental expression of ascidian neurotransmitter synthesis genes. Development Genes and Evolution 2002, 212(1):50–53.

44. Ikuta T, Saiga H: Dynamic change in the expression of developmental genes in the ascidian central nervous system: revisit to the tripartite model and the origin of the midbrain-hindbrain boundary region. Dev Biol 2007, 312.

45. Razy-Krajka F, Lam K, Wang W, Stolfi A, Joly M, Bonneau R, Christiaen L: Collier/OLF/EBF-dependent transcriptional dynamics control pharyngeal muscle specification from primed cardiopharyngeal progenitors. Developmental cell 2014, 29(3):263–276.

46. Imai JH, Meinertzhagen IA: Neurons of the ascidian larval nervous system in Ciona intestinalis: I. Central nervous system. Journal of Comparative Neurology 2007, 501(3):316–334.

47. Nishino A, Baba SA, Okamura Y: A mechanism for graded motor control encoded in the channel properties of the muscle ACh receptor. Proceedings of the National Academy of Sciences 2011, 108(6):2599–2604.

48. Kobayashi M, Takatori N, Nakajima Y, Kumano G, Nishida H, Saiga H: Spatial and temporal expression of two transcriptional isoforms of Lhx3, a LIM class homeobox gene, during embryogenesis of two phylogenetically remote ascidians, Halocynthia roretzi and Ciona intestinalis. Gene Expression Patterns 2010, 10(2):98–104.

49. Christiaen L, Stolfi A, Davidson B, Levine M: Spatio-temporal intersection of Lhx3 and Tbx6 defines the cardiac field through synergistic activation of Mesp. Developmental biology 2009, 328(2):552–560.

50. Takatori N, Kumano G, Saiga H, Nishida H: Segregation of Germ Layer Fates by Nuclear Migration-Dependent Localization of Not mRNA. Developmental Cell 2010, 19(4):589–598.

51. Tsagkogeorga G, Turon X, Hopcroft RR, Tilak M-K, Feldstein T, Shenkar N, Loya Y, Huchon D, Douzery EJ, Delsuc F: An updated 18S rRNA phylogeny of tunicates based on mixture and secondary structure models. BMC Evolutionary Biology 2009, 9(1):187.

52. Katsuyama Y, Okada T, Matsumoto J, Ohtsuka Y, Terashima T, Okamura Y: Early specification of ascidian larval motor neurons. Developmental biology 2005, 278(2):310–322.

53. Dynes JL, Ngai J: Pathfinding of olfactory neuron axons to stereotyped glomerular targets revealed by dynamic imaging in living zebrafish embryos. Neuron 1998, 20(6):1081–1091.

54. Kimmel CB, Model PG: Developmental studies of the Mauthner cell. In: Neurobiology of the Mauthner Cell. Raven Press New York; 1978: 183–220.

55. Denoeud F, Henriet S, Mungpakdee S, Aury J-M, Da Silva C, Brinkmann H, Mikhaleva J, Olsen LC, Jubin C, Cañestro C: Plasticity of animal genome architecture unmasked by rapid evolution of a pelagic tunicate. Science 2010, 330(6009):1381–1385.

56. Fujii S, Nishio T, Nishida H: Cleavage pattern, gastrulation, and neurulation in the appendicularian, Oikopleura dioica. Development genes and evolution 2008, 218(2):69–79.

57. Seo H-C, Kube M, Edvardsen RB, Jensen MF, Beck A, Spriet E, Gorsky G, Thompson EM, Lehrach H, Reinhardt R: Miniature genome in the marine chordate Oikopleura dioica. Science 2001, 294(5551):2506–2506.

58. Seo H-C, Edvardsen RB, Maeland AD, Bjordal M, Jensen MF, Hansen A, Flaat M, Weissenbach J, Lehrach H, Wincker P: Hox cluster disintegration with persistent anteroposterior order of expression in Oikopleura dioica. Nature 2004, 431(7004):67.

59. Edvardsen RB, Lerat E, Maeland AD, Flåt M, Tewari R, Jensen MF, Lehrach H, Reinhardt R, Seo H-C, Chourrout D: Hypervariable and highly divergent intron-exon organizations in the chordate Oikopleura dioica. Journal of molecular evolution 2004, 59(4):448–457.

60. Nishida H: Development of the appendicularian Oikopleura dioica: culture, genome, and cell lineages. Development, growth & differentiation 2008, 50(s1).

61. Conklin EG: The organization and cell-lineage of the ascidian egg, vol. 13; 1905.

62. Hotta K, Mitsuhara K, Takahashi H, Inaba K, Oka K, Gojobori T, Ikeo K: A web‐based interactive developmental table for the ascidian Ciona intestinalis, including 3D real‐image embryo reconstructions: I. From fertilized egg to hatching larva. Developmental dynamics 2007, 236(7):1790–1805.

